# QsRNA-seq: a method for high-throughput profiling and quantifying small RNAs

**DOI:** 10.1101/265603

**Authors:** Alla Fishman, Dean Light, Ayelet T. Lamm

## Abstract

The finding that small non-coding RNAs (sRNAs) can affect cellular processes by regulating gene expression had a significant impact on biological research and clinical diagnosis. Yet, the ability to quantify and profile sRNAs, specifically miRNAs, using high-throughput sequencing is especially challenging because of their small size and repetitive nature. We developed QsRNA-seq, a method for preparation of sRNA libraries for high-throughput sequencing that overcomes this difficulty by enabling separation of fragments shorter than 100nt long that differ only by 20nt in length. The method supports using unique molecular identifiers for quantification. We show that QsRNA-seq gives very accurate, comprehensive and reproducible results. Using QsRNA-seq to study the miRNA repertoire in *C. elegans* embryo and L4 larval developmental stages, enabled extending the list of miRNAs that are expressed in a developmental-specific manner. Interestingly, we found that miRNAs 23nt long are predominantly expressed in developmental stage L4, suggesting a possible connection between the length of miRNA and its developmental role.

## Introduction

Small RNAs (sRNAs) are 20-30 nucleotide long non-coding RNA molecules that impact diverse biological events through the control of gene expression and genome stability. During the last decade small RNAs emerged as central players in the regulation of gene expression in all kingdoms of life [1] and have been shown to regulate virtually all cellular processes. There are several classes of sRNA [2], with Dicer generated microRNAs (miRNAs) being the most extensively studied [3].

High-throughput sequencing (HTS) is currently the method of choice to identify and analyze the cellular repertoire of RNAs because of this method’s ability to investigate the entire transcriptome in an unbiased way. While preparation of mRNA and DNA libraries for HTS became a routine procedure, preparation of sRNA libraries remains technically challenging. Most protocols for generation of sRNA libraries require the sRNA molecules to be ligated from both sides (5’ and 3’) to oligonucleotide adapters that contain the sequencing primers, reverse transcribed for cDNA generation, and amplified by PCR [4]. The first challenge that arises at the very beginning of library preparation is separation of the sRNAs from other RNA species close in size, including tRNA and rRNA. Omission of this step results in sRNA libraries that are highly contaminated. In addition, the adapters readily ligate to each other instead of to the RNA, and this product (adapter-dimer), if not removed, is preferably amplified by the PCR, resulting in null sequences in the HTS data. Because of the small size difference between sRNAs and tRNA/rRNA and between the sRNA ligated product and adapter-dimer, it is very difficult to separate them, which affects the quality of the library. Denaturing polyacrylamide gel (PAGE), commonly used for size-based separation of small fragments, is not just time consuming and requires expertise, but also results in a significant loss of the product and eliminates the possibility of automatization. Solid Phase Reversible Immobilization (SPRI) on magnetic beads [5, 6] is widely used for nucleic acid separation based on size, however this method is not capable of discriminating between fragments shorter than 100 nucleotides and thus is not applicable for preparation of sRNA libraries.

The implementation of HTS for sRNA profiling is also hindered by the inability to reliably quantify the output data. Small RNA library construction for HTS involves a PCR amplification step, which is prone to bias. Because PCR is not a linear process, it quickly reaches plateau, distorting expressional differences. Moreover, PCR efficiency depends on the length of the fragment and on its sequence; variations in base composition might lead to preferred template-specific amplifications [7]. HTS-based miRNA expression data is thus not regarded as quantitative, and other techniques, mostly quantitative real-time PCR (qPCR) and miRNA microarrays, are used instead for quantification of individual miRNA and for large-scale studies, respectively. While these two assays are accurate and sensitive, both require prior knowledge and accurate annotation of the miRNA sequence tested and are not applicable to discovering novel miRNAs.

PCR-derived artifacts can be corrected by counting absolute numbers of molecules using Unique Molecular Identifiers (UMI) [8]. This method allows distinguishing between original copies of the sRNA present in cells and their amplification products by marking, prior to the amplification step, each molecule in a population by attaching a UMI, a short random sequence. Following amplification, each one of the UMIs attached to an original copy of the molecule will be observed multiple times; however, the original copy number of a molecule can be determined simply by counting each UMI only once upon analysis of HTS sequencing data. Thus, UMIs in the library acts as a molecular memory for the number of molecules in the starting sample. Surprisingly, this method is rarely used for generation of sRNA libraries. Here, we present QsRNA-seq, a novel method for preparation of small RNA libraries for HTS sequencing that overcomes the abovementioned shortcomings. Our protocol comprises of two innovations: (1) Gel-less size based separation of fragments shorter than 100nt, differing in length in 20 nt or more. (2) Utilization of UMI, enabling quantification of small RNA expression data.

## Results

### A novel method for separating nucleic acids shorter than 100 nucleotides

Most of the difficulties in sequencing and quantifying small RNAs derive from their small size and repetitive nature. To single them out and separate them from rRNA and tRNA and to separate ligation products from adapter-dimers, a technique that will allow a good and simple separation based on size of fragments ranging between 20 to 100 nucleotides is required. To achieve this, we modified SPRI [5, 6], a size-selection method based on a non-specific reversible binding of nucleic acid molecules to carboxyl groups coated magnetic beads in the presence of a “crowding agent” such as polyethylene glycol (PEG). As the efficiency of the binding is dependent on the length of the fragment and the concentration of the crowding agent, it is possible to separate two fragments of different lengths. It is long known that an addition of Isopropanol, another crowding agent, to PEG, modifies the range of bound fragment sizes, allowing binding of molecules to the magnetic beads as short as 18nt. Our hypothesis was that by adjusting the concentration of Isopropanol added to PEG, we would be able to achieve separation of molecules shorter than the 100nt threshold [9]. Therefore, we prepared a series of SPRI based size-selection solutions, all having the same concentration of PEG (7.5%) but different concentrations of Isopropanol, ranging from 32% to 54.5%, and tested their ability to promote binding of synthetic single stranded DNA oligonucleotides of different lengths to the beads (see methods). The oligonucleotides sizes, ranging from 19 to 66nt, were chosen to cover the separation steps needed for small RNA library preparation, namely separation of 3’adapter ligated sRNA from free 3’adapter and separation of 3’,5’- adapter ligated sRNA from adapter dimer. Binding efficiency was calculated by the ratio of oligonucleotide quantities in the eluent versus the input using a flurometer. The results of the experiment are summarized in Table 1. As hypothesized, increasing concentration of Isopropanol leads to an increase in the binding efficiency. Moreover, for each oligo length tested, we determined a condition resulting in its significant binding (>40%) to the beads, while oligos shorter by 20 nucleotides bound poorly (<5%). We next tested the feasibility of using these conditions to separate two oligonucleotides, 37nt and 58nt, differing in length by 21nt. We used double-sided size-selection on the SPRI beads, by 1. binding the longer fragment to the beads and collecting the unbound material containing the shorter fragments (right-side size-selection) 2. adding a second batch of beads, and adjusting the conditions (based on Table 1) to allow complete binding of the shorter fragments (left-side size-selection). Eluting the beads from the right-size selection will isolate the longer fragments and eluting the beads from the left size selection will provide the shorter fragments. For isolating the 58nt oligos, we used 3 different concentrations of Isopropanol for the right-side size selection, 38%, 41%, and 44%. The supernatant, containing the unbound shorter oligonucleotide, was transferred to a new tube and a left-size selection was performed at 54.5% Isopropanol to allow maximal recovery of the 37nt oligos. The input mixture and eluates from each size-selection step were analyzed using Tapestation (Figure 1). Binding efficiencies were consistent with those determined using a single oligo (Table 1). Using 38% Isopropanol at the right-size selection recovered around 2/3 of the 58nt input material with minor left overs of the 37nt oligos (Figure 1A,B), while the left-size selection resulted in nearly a complete recovery of the 37 oligos but a third of the input material of the 58nt oligos (Figure 1C). A mirror picture was obtained using 44 % Isopropanol, right-size selection resulted in a complete recovery of the 58nt oligos with a noticeable fraction of 37nt oligos (Figure 1G,H), while the left-size size selection yielded third of the 37nt input oligos with almost no 58nt oligos (Figure 1I). The in-between results were observed when using 41% Isopropanol (Figure 1 D,E,F). We concluded that by using the concentrations of Isopropanol presented in Table 1, it is possible to separate between two short nucleic acids differing by even less than 20nt with high recovery.

**Table 1.** Binding efficiencies of ssDNA oligonucleotides to SPRI beads at variant Isopropanol concentrations. 100ng ssDNA oligonucleotides were brought to a total volume of 50ul with H2O and appropriate amount SPRI beads (in 20% PEG, 2.5M NaCl) and 100% Isopropanol were added to obtain the desired concentrations (for added quantities see methods). Oligonucleotides bound to beads were next separated, washed and eluted. Oligonucleotide quantities in the eluate and in the input solution were determined by fluorometer. Binding efficiency was calculated as the percentage of eluted ssDNA from the input quantity. Each data point represents an average of at least 3 separate experiments. ND-not determined.

| Oligo size<br>(nt) | Isopropanol concentration (%) |  |  |  |  |  |  |  |  |
| --- | --- | --- | --- | --- | --- | --- | --- | --- | --- |
|  | 30 | 32 | 35 | 38 | 41 | 44 | 48 | 51 | 54.5 |
| 66 | 21 | 56 | 74 | 87 | 97 | 100 | ND | ND | ND |
| 58 | ND | 4 | 8 | 42 | 80 | 89 | ND | ND | ND |
| 44 | ND | 2 | 3 | 13 | 40 | 59 | 75 | ND | ND |
| 37 | ND | ND | 1.5 | 4 | 11 | 19 | 45 | 80 | ND |
| 30 | ND | ND | ND | ND | 4 | 5 | 20 | 40 | 61 |
| 21 | ND | ND | ND | ND | ND | 2 | 4 | 7 | 15 |
| 19 | ND | ND | ND | ND | ND | 1 | 3 | 3 | 5 |

**Figure 1.**
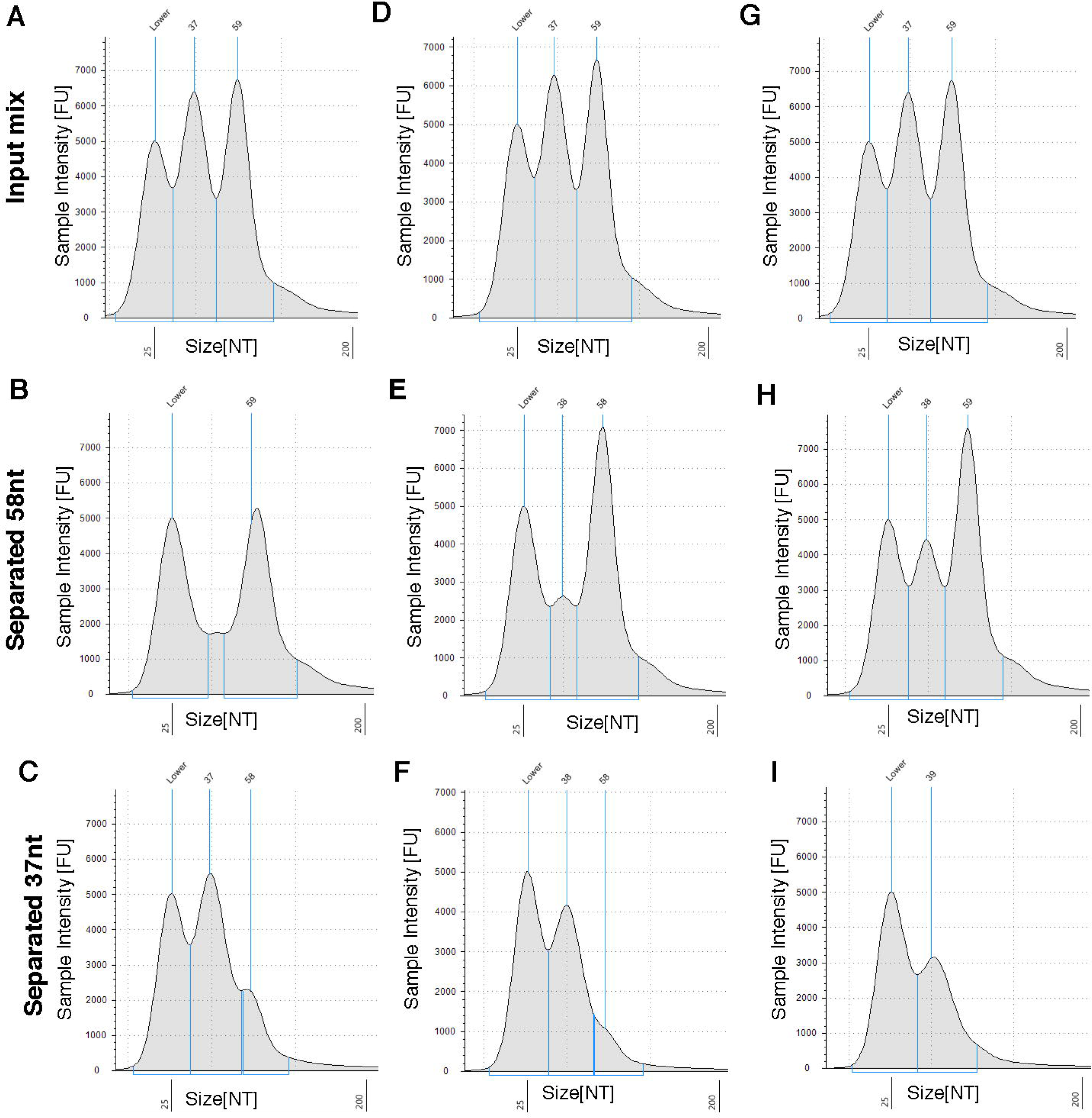
Separation of 37nt and 58nt fragments. Tapestation traces of an input mix of two ssDNA oligonucleotides, 37nt and 58nt, separated by double-sided size- selection on SPRI beads. The right-side size-selection was performed at three different Isopropanol conditions, 38% (A,B,C), 41% (D,E,F), and 44% (G,H,I). Input oligonucleotide mixtures for each concentration are presented in (A,D,G). Eluates of the right-side size selection using each Isopropanol concentration are presented in (B,E,H). Eluates of the left-side size selection using each Isopropanol concentration are presented in (C,F,I). Peak sizes and corresponding fragments areas are marked in blue; the left peak titled “lower” is a 25nt size marker.

### QsRNA-seq - a method for preparation of small RNA libraries

We next designed a new protocol for preparation of small RNA libraries for high-throughput sequencing, utilizing the separation method we developed. The protocol, named QsRNA-seq, is presented in Figure 2A (for detailed protocol see Supp protocol). The protocol, based on [10], [11], implements two ligation steps: 1. ligation of pre-adenylated 3’-adapter without ATP and 2. ligation of 5’-adapter containing a 4-nt barcode to allow multiplexing. Three size-separation steps on SPRI magnetic beads are performed during the protocol to obtain only the required RNA molecules: 1. separation of small RNA from longer RNAs (being mainly tRNA) prior to the first ligation. 2. separation of 3’-adapter ligated small RNA from free 3’- adapter following the first ligation. 3. separation of 3’,5’-adapter ligated small RNA from adapter-dimer and free 5’-adapter following the second ligation (for sizes of fragments see Suppl. Table 1).

**Figure 2.**
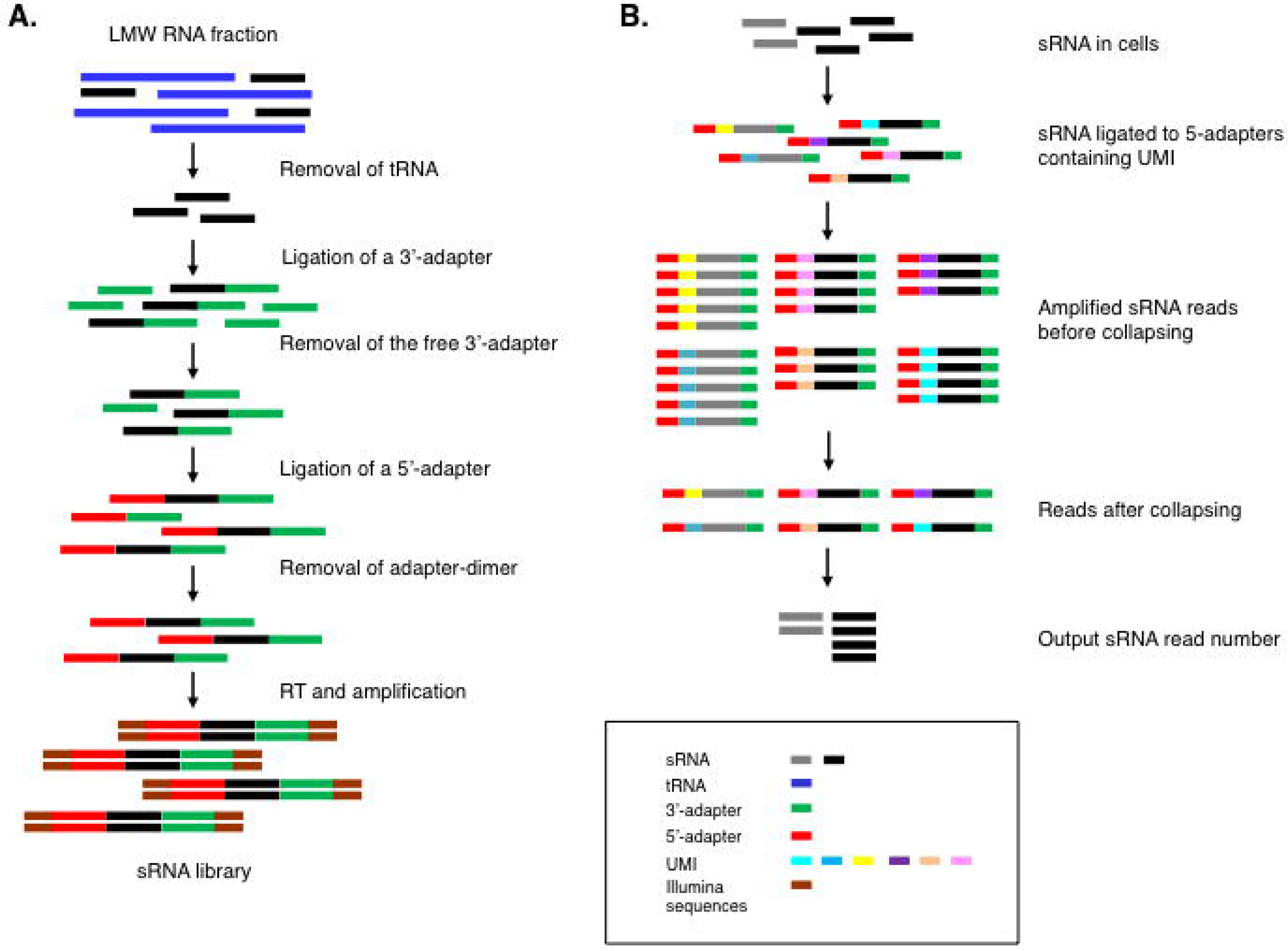
QsRNA-seq library preparation scheme. A. a general scheme for preparation of sRNA library for high-throughput sequencing from low molecular weight (LMW) RNA fraction. sRNA is separated from tRNA and then ligated to 3’-adapter. Next, the 3’-ligated sRNA is separated from the remaining free 3’-adapter and then ligated to 5’-adapter possessing UMI. The 3’-5’ ligated sRNA is separated from free 5’-adapter and from 3’-5’- adapter-dimer and subjected to Reverse transcription (RT) and PCR amplification. All the separation steps of the protocol are performed by size-selection using SPRI magnetic beads. B. Schematic view of integrating Unique Molecular Identifiers (UMIs) to sRNA library preparation. The UMI, residing in the 5’-adapter, enables marking each adapter-ligated miRNA with a unique random sequence before PCR amplification. Thus, after PCR amplification it is possible to distinguish identical sequences originating from the same molecule (having the same UMI) from sequences originating from different molecules.

In order to correct for PCR-induced artifacts and enable quantification, we used 5’ adapters that contain 8 random nucleotides that provide a unique identifier to each RNA molecule (UMI) [8]. After PCR amplification, we considered identical small RNA with the same UMI as an amplification product and merged them to one sequence (i.e., collapsing, Figure 2B).

To test the ability of QsRNA-seq to detect sRNAs, we used QsRNA-seq on RNA extracted from wildtype *C. elegans* synchronized to embryo or L4 larval stage and on total RNA obtained from human brain. Human brain total RNA was chosen because miRNAs constitute most of the small RNAs in this sample, thus we expected it to result in a very uniform library. In contrast, *C. elegans* contains many types of small RNA, including miRNAs, primary and secondary endogenous siRNAs, and piRNAs. We generated 3 independent biological samples from each *C. elegans* developmental stage, embryo and L4, for biological replica. RNA extracted from one sample from each stage was also subjected to 3 independent library preparations, as a technical replica, and was also used to prepare 3 technical replica libraries having no UMI in the 5’-adapter (0N). All library preparations resulted in very clean products ready for sequencing with negligible ratio of adapter-dimer containing no product (less than 2% of total reads in each library, examples in Supp. Figure 1, libraries information in Supp. Table 2).

### QsRNA-seq can evaluate miRNAs abundance and expression changes accurately

To evaluate the quality of the QsRNA-seq method output sequences, we aligned the generated sequences to all annotated miRNAs (both miRNA and miRNA*) in *C. elegans*, miRBase WBcel235, or in human, miRBase GRCh38. In the *C. elegans* sample, all the annotated miRNAs were present in our samples by at least one strand (3P or 5P), while 97% of all microRNAs, had coverage for both strands. Even rare miRNAs such as lsy-6, which is expressed in only one pair of neurons in the *C. elegans* head were present [12] (Supp Table 3). In addition, QsRNA-seq allows extensive multiplexing of the samples before amplification, which can reduce significantly the amount of starting material required. However, even without multiplexing, reducing the starting material by 10 fold, from 1ug to 100ng, produced nearly identical results (Supp. Figure 2). In human brain, the coverage was somewhat lower, with alignment to 80% of annotated miRNAs (Supp. Table 4). The difference probably derives from the large number of samples that we generated from whole worms at two developmental stages while we only generated one sample from human cells from a specific tissue. However miRNAs known to be enriched in human brain, for example, let-7 family, mir-9, mir-26a and others [13] were very abundant in our libraries.

To assess the consistency of the method, we evaluated the dispersion of miRNA expression between the replica samples, biological and technical, collapsed and non-collapsed. As expected, the collapsed replica exhibited lower dispersion rates than the corresponding non-collapsed replica, for both biological and technical replica types (Figure 3, Supp. Figure 3). For example, comparing the dispersion of the biological samples at embryo stage between collapsed reads and non-collapsed reads (Figure 3B,D), we observed at high normalized counts (> e+03) that the dispersion in the collapsed samples ranges between around e-0.5 and e-0.8, whereas the non-collapsed counts range between e-0.125 and e-0.22. Moreover, collapsed count dispersion decreases as the mean of the normalized counts increases, thus confirming our assumption that collapsing will tend to reduce statistical errors more drastically when dealing with larger counts.

**Figure 3.**
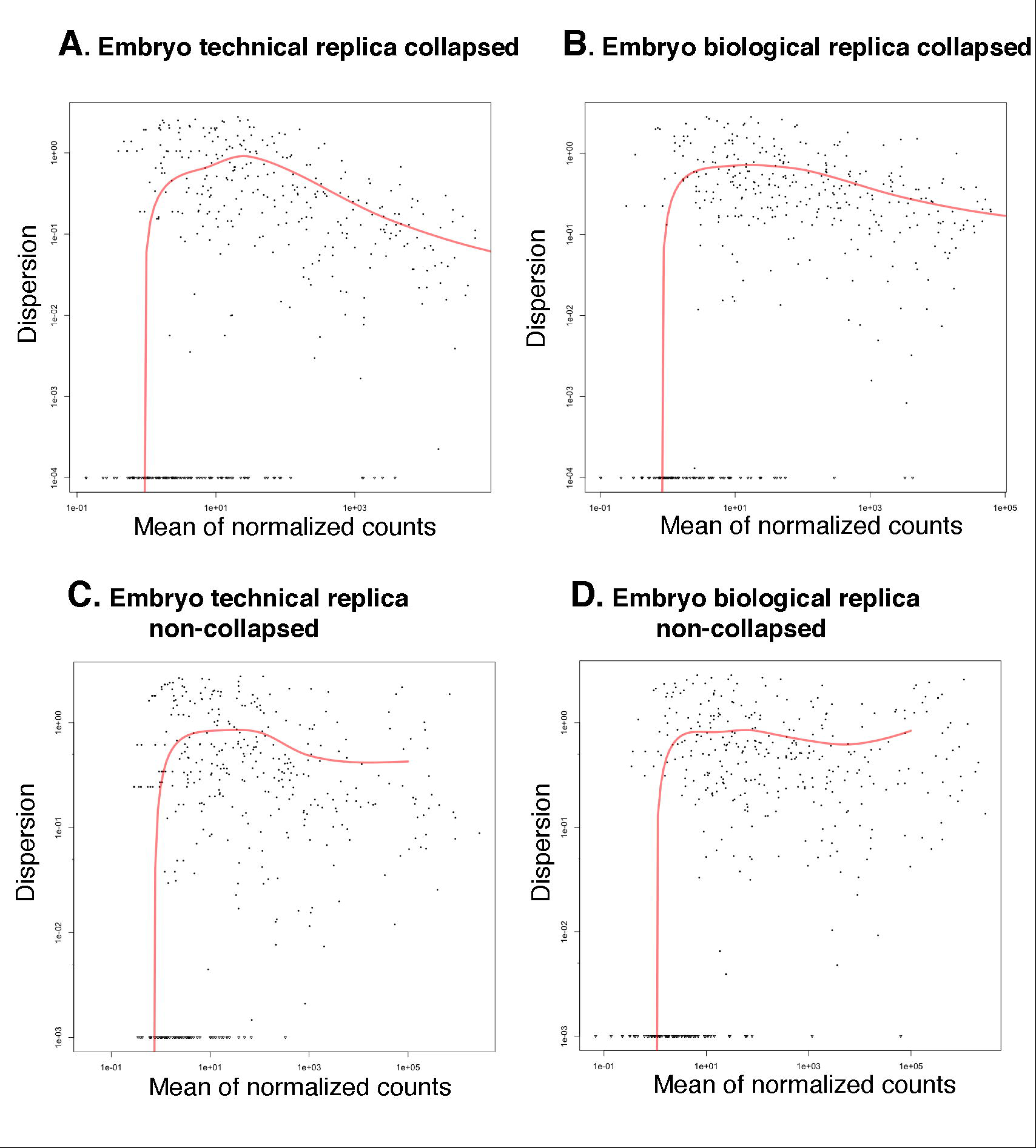
Variance between collapsed and non-collapsed biological and technical replica samples at embryo stage is low. Dispersion plots generated by DESEQ package in R using estimateDispersions function. Sequences aligned to each miRNA were counted and variance of samples was estimated. Each dot in the plot represents variance between samples for specific miRNA counts. Y-axis presents a dispersion value, which is the difference between samples squared. For example, a miRNA whose expression in different replicas differs by 10% will have a dispersion value of 0.01. X-axis is the mean of normalized counts. All plots are samples generated from embryo developmental stage, (A) technical replica samples dispersion estimated with collapsed reads, (B) biological replica samples dispersion estimated with collapsed reads, (C) technical replica samples dispersion estimated with non-collapsed reads, and (D) biological replica samples dispersion estimated with non-collapsed reads.

The increase in variation between samples, before and after collapsing, correlates with the abundance of the miRNA in the initial samples, due to a biased amplification of the abundant miRNA by PCR. While the ratio between the number of reads obtained for a single miRNA by collapsing and by non-collapsing is not significant for the low abundant miRNA (Suppl. Fig. 4A), it increases drastically in direct relation to miRNA abundance. Interestingly, the ratio becomes significant (above two-fold) when the initial expression level of a miRNA rises above 100 copies (Suppl. Fig.4A). Despite this, the higher read number obtained by non-collapsing does not affect the global picture of differential expression between L4 and embryo stages, i.e., whether a miRNA is upregulated or downregulated (Suppl. Fig. 5). However, non-collapsing augments the magnitude of L4/Embryo expressional fold change for a subset of miRNA (Suppl. Fig. 4B).

### 23-nt long small RNAs are enriched in L4 larvae and not in embryos in *C. elegans*

Besides miRNAs, which are mostly 22-nt long, *C. elegans* contains many other endogenously generated types of sRNAs [14]. Small interfering RNAs (siRNAs), which are very abundant in *C. elegans*, fall into two groups: 1. Primary siRNAs, which are 5’ monophosphate and are 26nt long. 2. Secondary siRNAs, which are 5’ triphosphate and are 21-22nt long. Another group are the equivalent of piRNAs in *C. elegans*, the 21U group, which are 21-nt long. To assess whether QsRNA-seq is capable of detecting all small RNA types, we performed length distribution on all the genome-aligned sequences from libraries generated from *C. elegans* embryo and L4 larva stages (Figure 4). As the libraries were prepared in a way that capture mostly RNAs with 5’ monophosphate and not 5’ triphosphate by direct ligation, we did not expect to have many secondary siRNAs in these samples. Surprisingly, we observed a major difference in sequence length distribution between samples generated from embryos and samples generated from L4 larva worms. We found that in embryos the most prominent length of sRNAs is 22-nt, while in L4 larva samples the prominent length is 23-nt (Figure 4). By removing sequences that aligned to miRNAs and performing a new length distribution we found that most of the 22-nt and 23-nt long sequences are miRNAs (Figure 4A,B black bars versus dark grey bars). Similar analysis performed on the 21-nt long sequences, showed that these sequences, detected in all the samples in considerable quantities, are mostly 21U (Figure 4A,B). We are expecting the 26-nt peak to be primary siRNAs. Sequences generated from technical replica or from samples without UMI showed similar length distribution (Supp Figure 6). As expected, length distribution performed on the human brain sample showed significant peaks at 21-23-nt long, composed mainly of miRNAs (Supp Figure 7).

**Figure 4.**
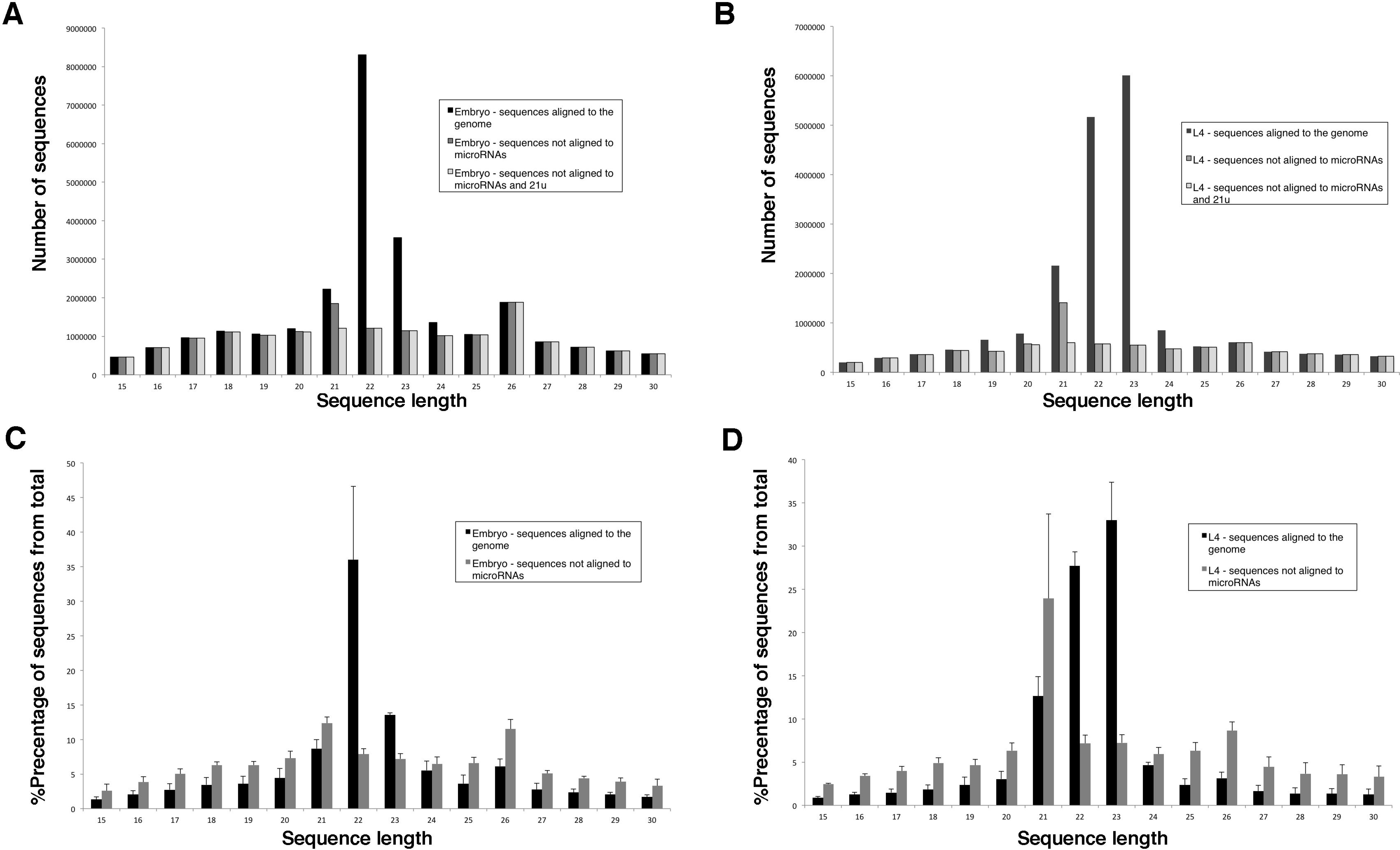
Size distribution of *C. elegans’s* sRNA sequences. (A,B) Bar charts presenting number of sequences for each sRNA sequence length from 15nt to 30nt. The sequences are from one sample from embryo stage (A) or L4 larval stage (B). (C,D) Bar graphs presenting percentage of sRNA for each sequence length from 15nt to 30nt from total number of sequences. (C) Average of 3 biological replicates at embryo stage. (D) Average of 3 biological replicates at L4 larval stage. Standard deviation is also presented. Black bars represent sequences that align to the genome, dark grey bars are sequences that did not align to miRNAs from sequences presented in the black bars, light grey bars represent sequences from the dark grey bar that also did not align to 21u sRNAs.

To further study the difference in length distribution of miRNAs between embryo and L4 samples, we evaluated the miRNA expression changes between embryo and L4 developmental stages and estimated the fold change difference. Selecting miRNAs with at least 5 fold difference and padj value < 0.05, we found 30 miRNAs that are predominantly expressed in embryo and 38 miRNAs that are predominantly expressed in L4 (Figure 5A, Supp Table 3). Our expression analysis is comparable to published data. For example, our findings that lin-4-family and let-7-family members are predominantly expressed during larval development, and miR-35 family members are predominantly expressed in embryogenesis, are in full concordance to data obtained using northern blot analysis [15, 16]. Our analysis is also comparable with large-scale studies performed either by multiplexed qPCR [17] (82% of predominantly expressed miRNAs in L4 and 100% in embryos were found by our analysis) or by HTS [18] (96% of predominantly expressed miRNAs in L4 and 55% in embryos found by our analysis). Our list of miRNAs differentially expressed between embryo and L4 stages was much longer than the corresponding lists of these two studies, mainly due to discovery of new miRNAs since these studies were performed. Our analysis was very restrictive, 3 biological replicas, 5 fold or more expression changes and padj value < 0.01, as compared to the other two analyses, which can explain the discrepancies between the lists.

**Figure 5.**
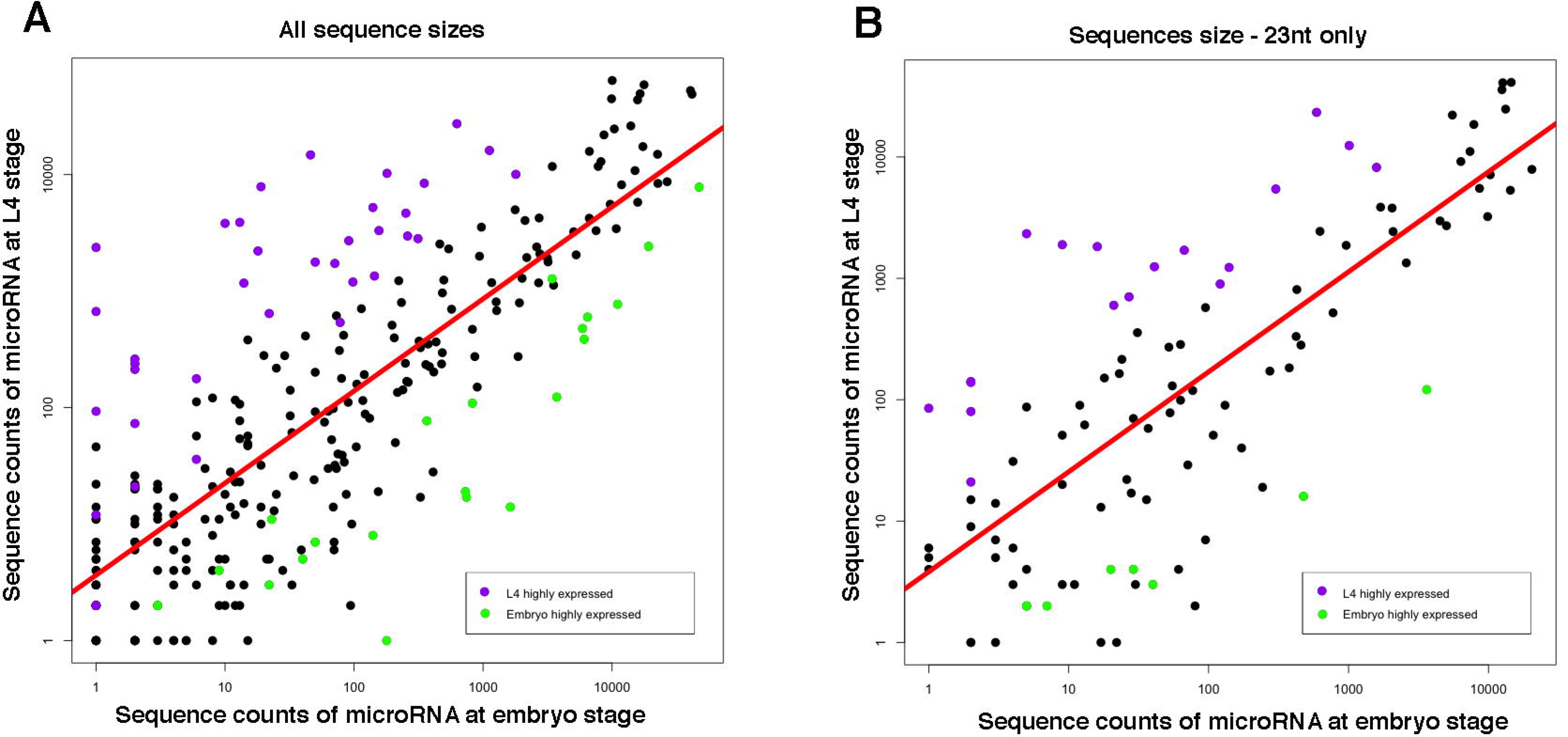
23nt long miRNAs are predominantly expressed in L4 larval stage. Log scale plot comparing miRNA expression between embryo and L4 developmental stages. A. All miRNAs B. 23nt long miRNAs only. Every dot in the plot represents a miRNA. Green dots indicate miRNAs predominantly expressed at embryo stage; purple dots indicate miRNAs predominantly expressed at L4 stage. The red line is the regression line for all miRNAs presented in the graph.

Interestingly, evaluating miRNA expression using only sequences that are 23nt long (Figure 5B), we found high fraction of 23nt long miRNAs that are predominantly expressed in L4 (45%) as compared to miRNA predominantly expressed in embryo stage (23%). Evaluating miRNA expression using sequences that are 22nt long, we did not observe a significant difference in the fraction of miRNAs predominantly expressed in L4 versus embryo stage (Supp Figure 8). Thus, we conclude that in *C. elegans* the expression of 23-nt long miRNAs is mostly L4 developmental stage specific.

## Discussion

Although sRNAs (20-30nt) constitute a very small fraction of the total cellular RNA, they have a significant role in every aspect of cellular and organismal development and maintenance [1]. Thus, identification, characterization, and quantification of sRNAs became an important part in many studies. miRNAs are also promising clinical diagnostic and predictive biomarkers [19]; in particular, miRNAs circulating in body fluids are attractive candidates to serve as markers for non-invasive “liquid biopsies” [20]. Alas, because of the difficulty in isolating sRNAs from other nucleic acids very close in size, profiling sRNAs using HTS is often avoided. To overcome this problem, we initiated a screen aimed to determine the best conditions of SPRI beads crowding agents to separate short nucleic acids fragments. This screen resulted in development of a method to separate fragments shorter than 100nt differing by as few as 20 nucleotides. The method also allows flexibility based on the trade off between purity and quantity (e.g. better separation between the fragments but less recovery of the desired fragment). While we concentrated on binding efficiency of fragments between 19 and 66nt, for needs of library preparation, binding conditions for fragments ranging between 60-100nt can be easily determined in the same manner for any other purpose.

By implementing the method for all the separation steps required for the preparation of high-quality small RNA libraries for HTS, we generated a new protocol, QsRNA-seq. The 20nt separation resolution that we gained is sufficient to significantly reduce the main two contaminants of sRNA libraries: tRNAs, and adapter-dimers. To further avoid adapter-dimers contamination, we also implemented in the protocol a common method of turning a free 3’- adapter into a double-stranded structure by hybridization the 3’-adapter with reversecomplimentary oligonucleotide [21]. Lack of contaminants resulted, as expected, in an impressive sequencing depth. Sequences obtained from two developmental stages of *C. elegans*, aligned to 97% of annotated *C. elegans* miRNAs. All other sRNA types known in *C elegans*, such as endogenous siRNA and 21U-RNA, were present in significant quantities. Degradation products similar in size to the small RNAs are probably also present in the sequencing data and cannot be avoided by this method, but can be minimized by using high-quality RNA.

While loss of material during QsRNA-seq is significantly reduced since no gel purification is required, an amplification step is still needed to produce enough material for HTS. PCR is usually used for amplification, however it is not a linear process and is not free of biases [7]. In contrast to the heterogeneity in reads observed in mRNA-seq libraries, due to random fragmentation of the input mRNA, reads obtained from miRNA libraries are very uniform, making quantification methods used in mRNA-seq, such as collapsing reads or FPKM, unsuitable for miRNA quantification. To allow quantification of miRNAs we used UMI to mark each molecule before the amplification step. Using UMI, we could collapse the sequencing reads similarly to what is done for mRNA-seq [22], obtaining quantitative global sRNA expression data. Another benefit of using UMI is that it allows unlimited number of PCR amplification cycles, enabling library preparation from low amount of starting material, which is especially beneficial for clinical purposes. UMI reliably reflects molecule counts only if the number of distinct labels is substantially larger than the copy number of the most abundant target molecule; copy number/labels ratio greater than 0.2 results in an approximate 10-25% undercounting of collapsed reads [23]. In our case, usage of 8-nt long UMIs resulted in random barcode pool being saturated for a fraction of highly overexpressed molecules (Supp Figure 4). Thus, when highly expressed miRNAs quantification is needed or low amount of starting material that requires many rounds of amplification is used, longer UMIs are preferable. But even with 8-nt long UMIs, reducing the input material by 10 fold produced very similar results (Supp Figure 2). QsRNA-seq can be adjusted easily to longer UMIs using Table 1.

The most surprising finding of our study is an expressional bias of 23-nt long miRNAs towards L4 larval stage in *C. elegans*, suggesting a possible connection between the length of miRNA and its role in the organism development. Interestingly, a study on endogenous siRNAs in *C. elegans* showed that targets of 23-nt siRNAs are associated uniquely with postembryonic development [24]. Generation of both endogenous siRNAs and miRNAs is depended on DICER enzyme and the competition among these processes on resources was suggested to affect development [25]. It is tempting to speculate that sRNA processing by DICER changes as development progress, which might be needed for normal development. Studies combining mRNA-seq and small-RNA sequencing might be able to shed more light on the function of these 23-nt long sRNAs.

## Conclusions

After established a way to separate between very short nucleic acid fragments (shorter than 100nt) that differ in length by only 20nt, we developed a new method, QsRNA-seq, to prepare small RNA libraries for HTS. QsRNA-seq is a gel-free, fast and easy to perform method that also utilizes UMI and barcoding, producing high quality sRNA libraries, generating high-depth expression data. We show that QsRNA-seq will be very useful for studies of small RNA data and for clinical diagnosis. In addition, profiling miRNA in *C. elegans* using QsRNA-seq, suggested that not just miRNA expression differs in different developmental stages but also miRNA sizes. We believe that QsRNA-seq can transform the preparation of small RNA libraries into a routine procedure as is the preparation of mRNA libraries.

## Methods

### *C. elegans* growth and synchronization

Wildtype *C. elegans* strain, Bristol N2, was used in this study and was maintained on OP50- seeded enriched plates at 20°C as described in [26]. Embryos were isolated from gravid N2 adults by treatment with sodium hypochlorite solution to dissolve animals of all stages but embryos. To obtain synchronized L4 worms, embryos were incubated in M9 media without food at 20°C for 24h. Hutched synchronized L1 were grown on OP50-seeded Enriched plates at 20°C until they reached L4 larval stage.

## RNA extraction

Synchronized embryos or L4 larval worms were washed several times with M9 to avoid contamination with bacteria, snap-freezed in liquid nitrogen and then grinded to powder by a liquid nitrogen pre-chilled mortar and pestle. High-molecular weight and low-molecular weight RNA fractions were isolated using miRVana miRNA isolation kit (Ambion). RNA quantity was measured by Qubit^®^ Fluorometer using Qubit^®^ RNA HS Assay Kit (Molecular probes) and its quality was estimated by agarose gel electrophoresis and Tapestastion (Agilent genomics). Human Brain RNA was obtained from (FirstChoiceuman brain Total RNA, Life Technologies).

## Determining SPRI binding conditions

Volumes of PEG solution and Isopropanol were calculated using the equation:

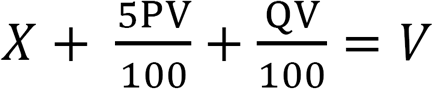

Where: V is total volume; X is volume of nucleic acid solution; P is desired concentration (%) of PEG; Q is desired concentration (%) of Isopropanol.

First, a total volume of binding solution (V) was calculated by substituting P, Q and X in the equation for the desired concentrations of PEG and Isopropanol and the volume of nucleic acid solution. Next, the volumes of 20% PEG and 100% Isopropanol needed for the desired concentrations, being equal to 5PV/100 and QV/100, respectively, were calculated.

To measure binding efficiency a solution of 2ng/ul of synthetic single-stranded DNA oligonucleotide was aliquoted 50 ul per tube. To each tube SPRI beads in 20% PEG (SPRIselect, Beckman-Coulter) and 100% Isopropanol were added at volumes determined using the calculation method above. Size-selection was performed according to the manufacturer’s protocol (Beckman’s AMpureXP, left-side selection). Oligonucleotide concentrations in the input and eluted samples were measured by Qubit^®^ Fluorometer using Qubit^®^ ssDNA Assay Kit (Molecular probes). Binding efficiency was calculated by the percentage of the output oligonucleotide from the input quantity.

## Small RNA library preparation

Small RNA libraries were prepared from at least 3 biological replicas of N2 worms at embryo or L4 stage. One RNA sample from each stage was selected for preparing two additional libraries, resulting in 3 technical replicas for each stage.

A step-by-step protocol developed in this study that includes reagents and primers can be found in the supplementary. In short, sRNA was separated from other RNA species and then ligated to 5’-adenylated 3’-adapter using T4 RNA ligase-2 truncated (NEB) in an absence of ATP. The 3’-adapter-ligated sRNA was separated from free 3’-adapters and then ligated to a 5’-adapter, containing multiplexing barcode and UMI, using T4 RNA ligase 1 (NEB). sRNA ligated from both sides was then separated from the adapter-dimer to obtain an sRNA library. All the separation steps of the process of library preparation were performed using the method described above of SPRI-based size-selection of short fragments. sRNA library was reverse-transcribed using QScript Flex cDNA synthesis kit (Quanta) and amplified using Phusion High-Fidelity DNA Polymerase (NEB). The amplified library was cleaned from primers and irrelevant products below 100bp and above 200bp by double-side size-selection on SPRI beads (Beckman’s AMpureXP) and its concentration and quality was determined by Tapestation analysis (Agilent genomics). Libraries were sequenced using 50 bp SR sequencing mode on HiSeq 2500 platform (Illumina).

## Sequences processing and expression analysis

RNA sequences obtained were first de-multiplexed according to the 4nt barcode. Next, the 3’adapter sequences were trimmed off by scanning from the 3’-end of the sequence the first instance of the adapter sequence by increments of 1nt. We then either (1) removed the barcode and UMI (8-nucleotide) and these sequences are considered as non-collapsed or (2) merged identical sequences and then removed the barcode and UMI and these sequences are considered as collapsed.

*C. elegans* sequences were either aligned to the WS220 (Wormbase, www.wormbase.org) genome using Bowtie [27] for size distribution analysis, allowing no mismatches with no more than 10 alignments to the genome or aligned to miRBase WBcel235 (www.mirbase.org), allowing no mismatches and not more then one alignment. Human brain sequences were aligned to miRBase GRCh38 with the same parameters.

Size distribution analysis was done on processed sequences before and after alignment to the genome. DEseq [28] package in R (http://www.r-project.org) was used to evaluate miRNA expression and estimateDispersions function in DEseq was used to estimate the dispersion between biological replicas and technical replicas.

## Declarations

### Funding

This work was funded by The Israeli Centers of Research Excellence (I-CORE) program, (Center No. 1796/12 to ATL), The Israel Science Foundation (grant No. 644/13 to ATL), and Israel Cancer Research Fund (ICRF).

### Availability of data and materials

All data generated in this study was submitted to NCBI/Genbank short read archive. A detailed protocol of QsRNA-seq can be found in the supplementary.

### Authors’ contributions

AF and ATL designed the experiments and protocol. AF performed the experiments. DL and ATL performed the bioinformatics analysis. AF and ATL wrote the manuscript.

### Ethics approval and consent to participate

Not applicable

### Competing interests

USA provisional patent application no 62/359,245 titled “Methods For Separation Of Short Nucleic Acid Molecules And Quantification Of Small RNA”, by A.F. and A.T.L, was submitted by Technion Research and Developmental Foundation on 07.07.2016

## Acknowledgements

We thank prof. Vladimir Lin, from the Technion Faculty of Mathematics, for his help in development of the equation for calculating volumes of PEG and Isopropanol. We also thank Orna Ben-Naim Zgayer, Nabeel Ganem, and Noa Ben-Asher for critical reading of the manuscript and discussions.

